## Supplementary figures and references for "Localized and global representation of prior value, sensory evidence, and choice in male mouse cerebral cortex"

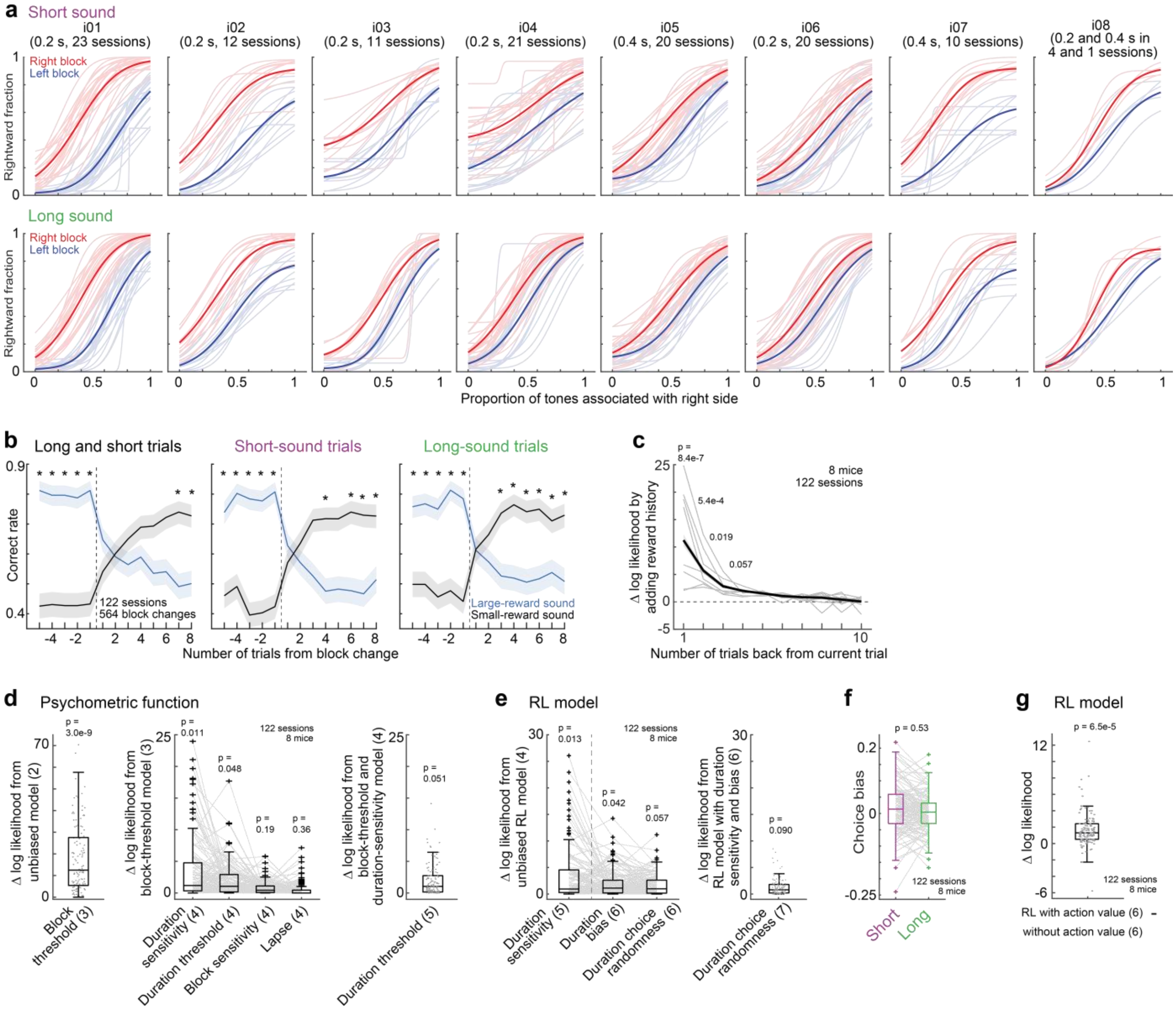


**Supplementary Fig. 1. Choice behavior in tone frequency discrimination task.**
**a.** Psychometric function of choice behavior. Bold lines show the average psychometric function in each mouse. Overlayed traces show the psychometric function in each session. Parentheses show the duration of short sound and the number of sessions in each mouse.

**b.** Correct-response rate for moderate and difficult tone clouds around block changes. Blue and black lines show the correct rate associated with large and small rewards in previous blocks, respectively. Means and 95% confidence intervals (*, $p < 1e-4$ in the two-sided t-test in linear mixed-effects model; 564 block changes from 122 sessions, 8 mice).

**c.** Reward history affected the choice in current trial. Regression analysis quantified how the outcomes of past trials in left and right choice were correlated to the current choice (**Methods**). The two-sided likelihood ratio test was used to investigate how the additional number of trials back from current trial significantly increased the prediction accuracy of choice (Δ log likelihood). Means and standard errors (122 sessions, 8 mice). Overlayed trace shows the average Δ log likelihood in each mouse. We found that the reward history of multiple trials affected the current choice.

**d.** Behavioral model comparison in psychometric function. We first analyzed the average log likelihood per session in each mouse, and then averaged across mice for the two-sided likelihood ratio test (**Methods**). Parentheses show the number of parameters in the psychometric-function models. The unbiased model had the parameters for stimulus sensitivity and choice threshold irrespective of blocks and sound durations (2 parameters). The two-sided likelihood ratio test was used to investigate whether the model with additional block-dependent or duration-dependent parameters fit mice choices ($p < 0.05$). The model with block-dependent threshold and duration-dependent sensitivity fit the mice choices (4 parameters) (central mark in box: median; edges of the box: 25th and 75th percentiles; whiskers: most extreme data points not considered outliers (beyond 1.5 times the interquartile range), here and throughout).

**e.** Model comparison in reinforcement learning (RL) model. Data are presented as in **d**. The unbiased RL model had the parameters of learning rate, choice randomness (inverse temperature), stimulus sensitivity, and choice-threshold irrespective of blocks and sound durations (4 parameters). The two-sided likelihood ratio test was used to investigate whether the duration-dependent sensitivity, threshold, and choice randomness improved the model fitting ($p < 0.05$). The RL model with duration-dependent sensitivity and choice bias fit the mice choices (6 parameters).

**f.** Duration-dependent choice bias in the RL model. The choice bias did not have significant differences between the short and long sounds (two-sided t-test in linear mixed-effects model).

**g.** Comparison of RL models with and without action value. The RL model with action value fit the mice choices better than our previous model which only used prior value for value updating (two-sided t-test in linear mixed-effects model)[1]. Source data are provided as a Source Data file.

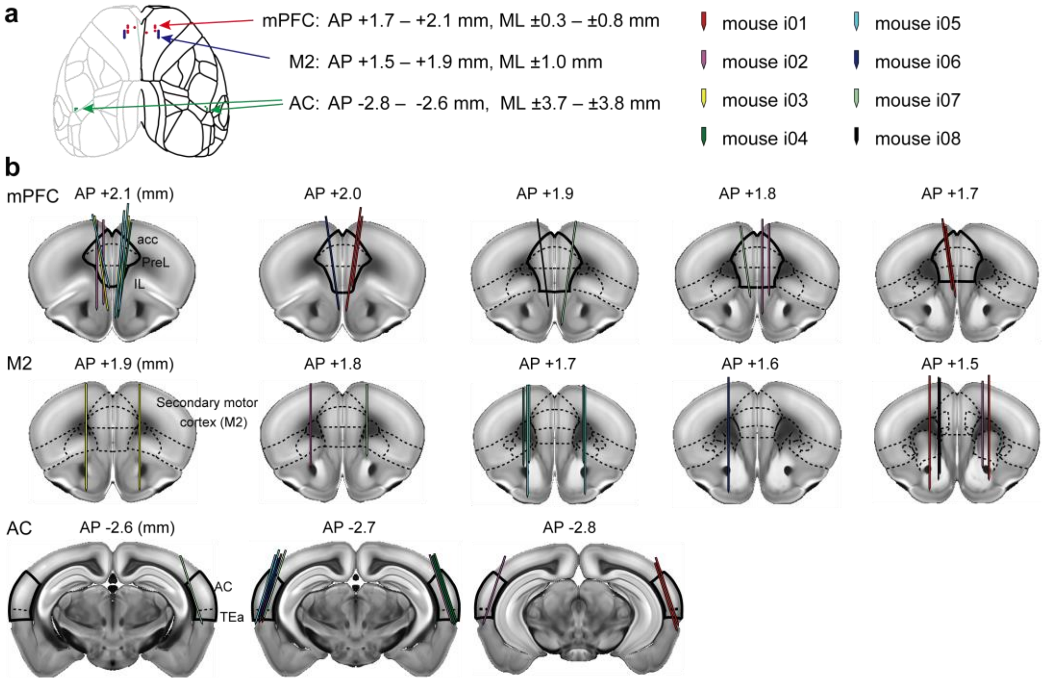


**Supplementary Fig. 2. Locations of Neuropixels probes**

**a**. Probe entry locations from the surface of dorsal cortex based on the Allen Brain Atlas coordinates. Red dots: the medial prefrontal cortex (mPFC). Blue dots: the secondary motor cortex (M2). Green dots: the auditory cortex (AC). AP and ML indicate the anterior-posterior and medial-lateral axes from bregma (mm), respectively. In each brain region and hemishere, we prepared one tiny hole in the skull for probe entry. In 34 out of 42 holes (80.9%), we identified the probe locations in the post hoc fixed brain slices (**Methods**). All the identified probe locations were successfully located in the target regions.

**b**. Probe tracks in 8 mice on coronal brain slices registered on Neuropixels trajectory explorer (https://github.com/petersaj/neuropixels_trajectory_explorer).

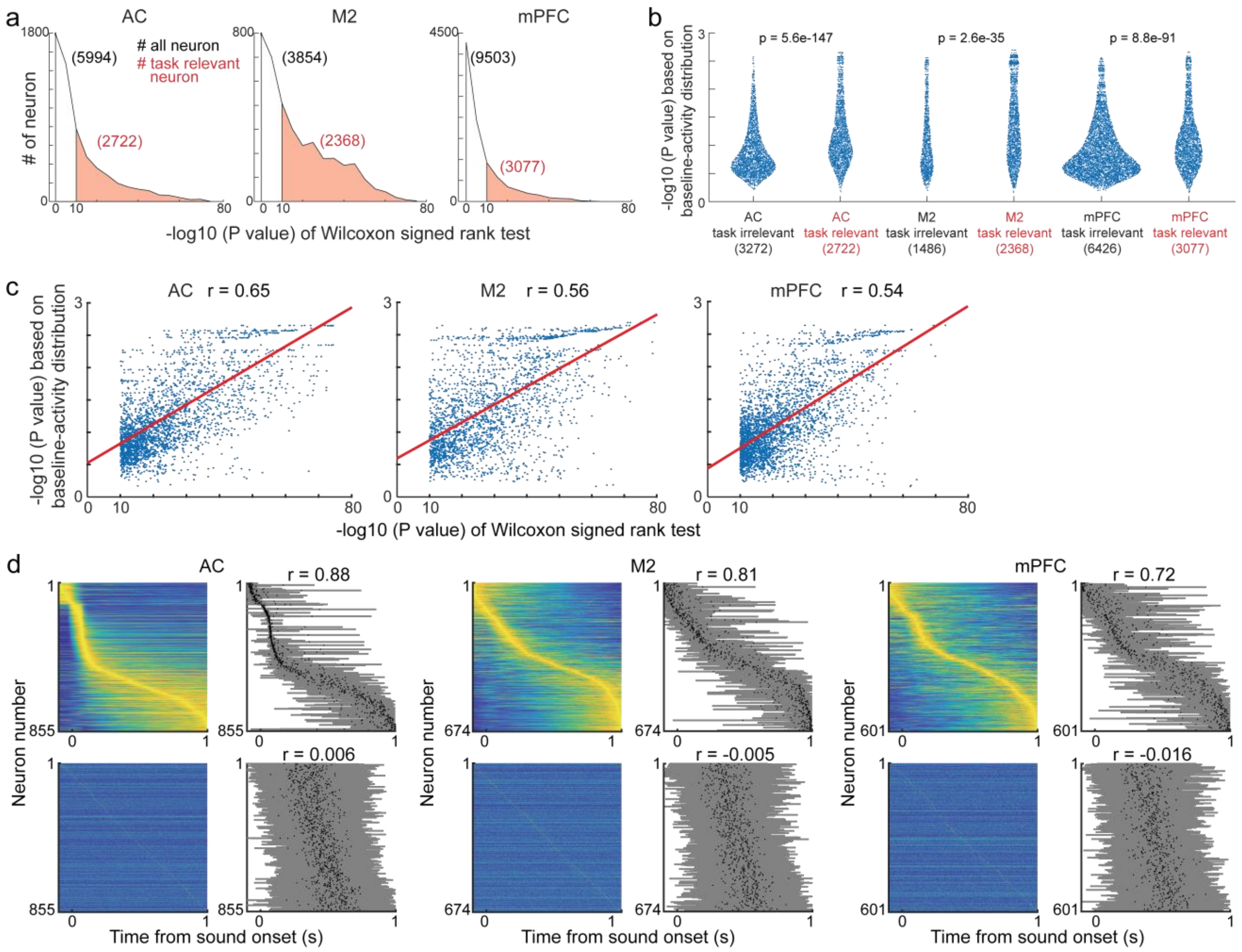


**Supplementary Fig. 3. Validation of task-relevant neurons**

**a.** Distribution of activity in all the recorded neurons. Task-relevant neurons (red areas) increased the activity during trial compared to the baseline activity during inter-trial-interval for sound-aligned activity or during before choice for choice-aligned activity (one-sided Wilcoxon signed rank test, p < 1e-10) (**Methods**).

**b.** Distribution of neural activity during trial compared with the baseline-activity distribution. The definition of baseline activity was identical to that for the task-relevant neurons in **a** (**Methods**). We first analyzed the distribution of baseline activity in each neuron. Based on the baseline distribution, we analyzed the probability (p value in y axis) of observing the activity during trial. When the activity during trial exceeded the baseline distribution, we set the inverse of the number of trials per session as the minimum p value. The task-relevant neurons defined by Wilcoxon signed rank test had smaller p values than the task-irrelevant neurons in all the three regions (two-sided Mann-Whitney U test).

**c**. Relationship between the analysis of Wilcoxon signed rank test (**a**) and that of the baseline-activity distribution (**b**) in task-relevant neurons. The p values of these two analyses were correlated (Spearman correlation). Red line shows the robust linear regression.

**d.** Distribution of neural activity in long sound trials (-0.1 - 1.0 s from the sound onset). Same as **Fig. 4a, 5f and Supplementary Fig. 9f**, a subset of task-relevant neurons which increased the activity between -1.0 and 1.0 s from sound onset are shown. (Left) In each neuron, we first applied a Gaussian filter with the standard deviation of 100 ms to the spike activity. We then averaged the filtered activity across trials and scaled between 0 and 1, sorted by the maximum activity timings. Bottom panels shuffled the activity of each neuron in time and analyzed the maximum activity timings. The average spike rates of each neuron were preserved. (Right) Stability of temporal activity patterns. First, we randomly split the long-sound trials into half. In each neuron, we averaged the activity during the half of trials and analyzed the maximum activity timing. The maximum activity timings of neurons were also analyzed in the other half of trials. Spearman correlation was analyzed between the maximum activity timings of neurons in the two distinct trials. We repeated the procedure 100 times to reduce noise from random grouping of trials. The temporal orders of activity analyzed from two distinct trials were highly correlated ($r > 0.72$ on average from 100 comparisons) compared to those from the shuffled activity ($r < 0.0072$), suggesting the stable temporal patterns of activity in the AC, M2, and mPFC. Black dots and gray areas show the means and standard deviations of maximum activity timings. Source data are provided as a Source Data file.

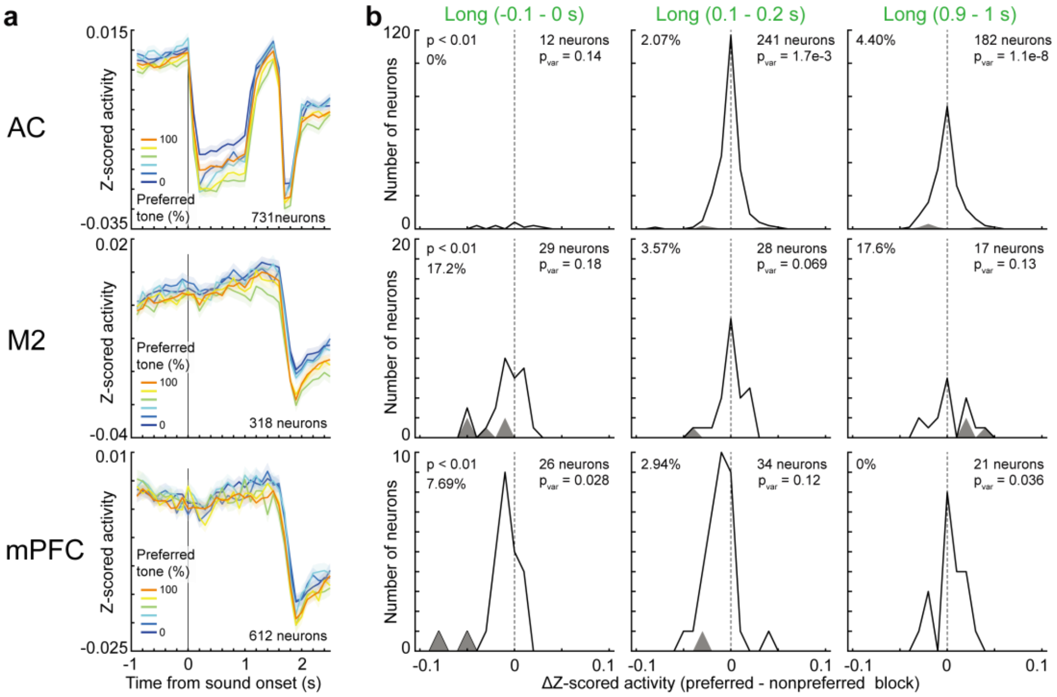


**Supplementary Fig. 4. Neurons decreasing the activity during trial**

**a.** We identified the neurons with significantly decreasing the activity during trial compared with the baseline activity. The definition of baseline activity was identical to that of task-relevant neurons (**Methods**) ($p < 1e\text{-}10$ in the one-sided Wilcoxon signed rank test). In the M2 and mPFC, the neurons mainly decreased the activity after mice made choices. Medians and robust standard errors.

**b.** Block modulations of the decreasing neurons in correct long-sound trials. We extracted the decreasing neurons in each time window and analyzed the difference in activity between the preferred and nonpreferred blocks. The gray areas show the neurons with significant change in activity between blocks ($p < 0.01$ in the two-sided Wilcoxon signed rank test). The proportion of significant neurons are shown as the figure insets. In the M2 and mPFC, the number of decreasing neurons were at least 10 times smaller than that of the increasing neurons in **Supplementary Fig. 5c**. In the AC, the decreasing neurons had smaller variance of block modulations than the increasing neurons ($P_{var}$ in the two-sided Levene's test). The distributions of increasing neurons are shown in **Supplementary Fig. 5c**. Source data are provided as a Source Data file.

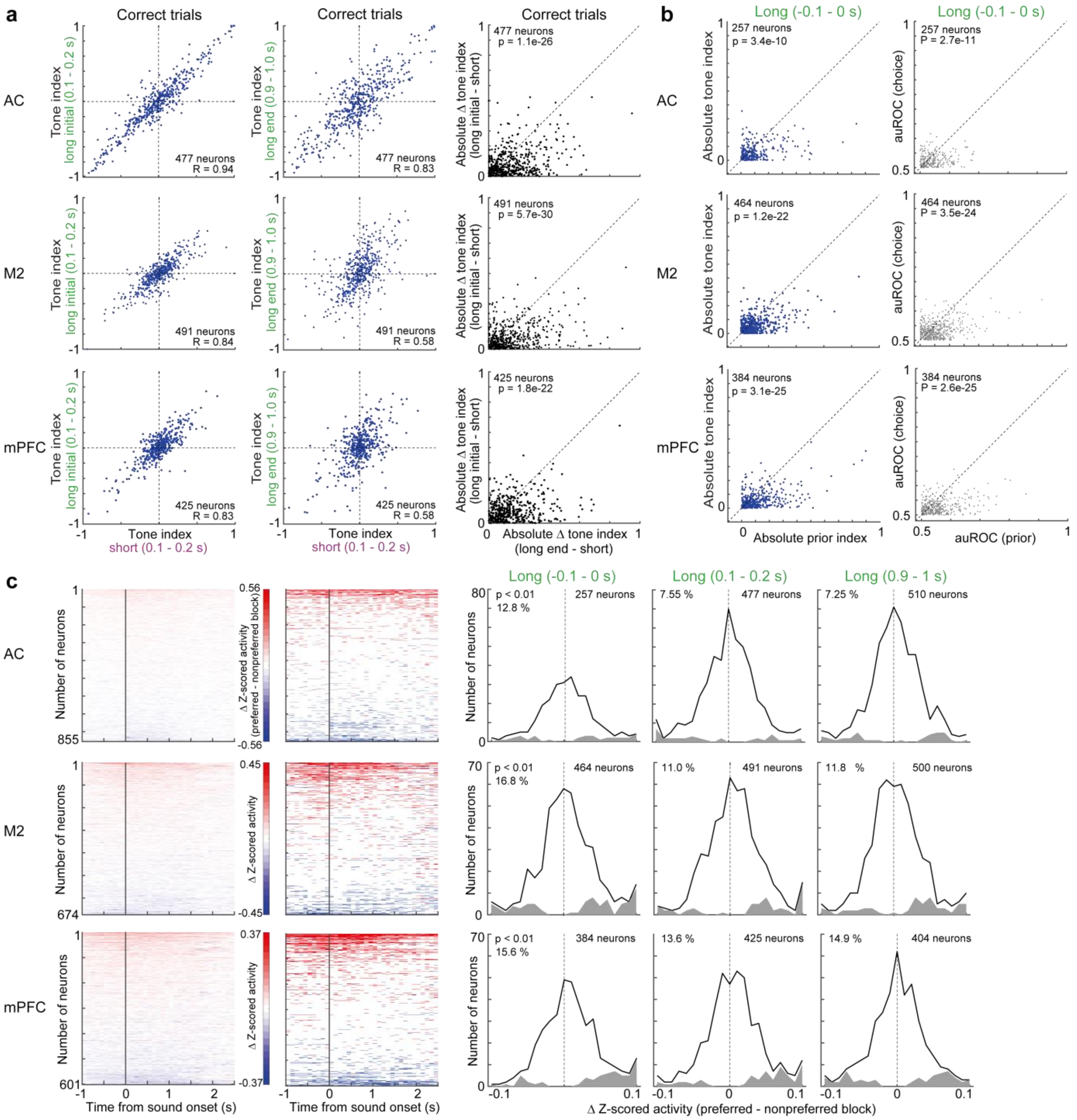


**Supplementary Fig. 5. Tone and prior indices during short- and long-sound trials.**

**a.** Tone index. We compared the tone indices between correct short- and long-sound trials. The tone indices of short-sound trials (0.1–0.2 s) were correlated to those of the initial long sounds (0.1–0.2 s) rather than to the end of long sounds (0.9–1.0 s) (left and middle: Pearson correlation; right: p value in the two-sided Wilcoxon signed rank test).

**b.** Prior index. The prior index was defined as the activity difference between the trials with left- and right-prior-value dominant trials. The absolute prior indices were larger than the tone indices before sound onset (left, two-sided Wilcoxon signed rank test). The performance of prior discrimination in single neurons were investigated with the area under the receiver operating curve (auROC) (**Methods**). The prior discrimination was better than the choice discrimination in all 3 regions (right, two-sided Wilcoxon signed rank test).

**c.** Block modulations of single neurons in correct long-sound trials. We analyzed the difference in activity between the preferred and nonpreferred blocks in each neuron (left). The time windows with significant block differences were colored as red (preferred) and blue (nonpreferred), respectively ($p < 0.01$ in the two-way ANOVA) (middle). In the right three panels, the gray areas show the neurons with significant change in activity between blocks ($p < 0.01$ in the two-sided Wilcoxon signed rank test). All 3 cortical regions had block-modulated neurons before and during long sounds. Source data are provided as a Source Data file.

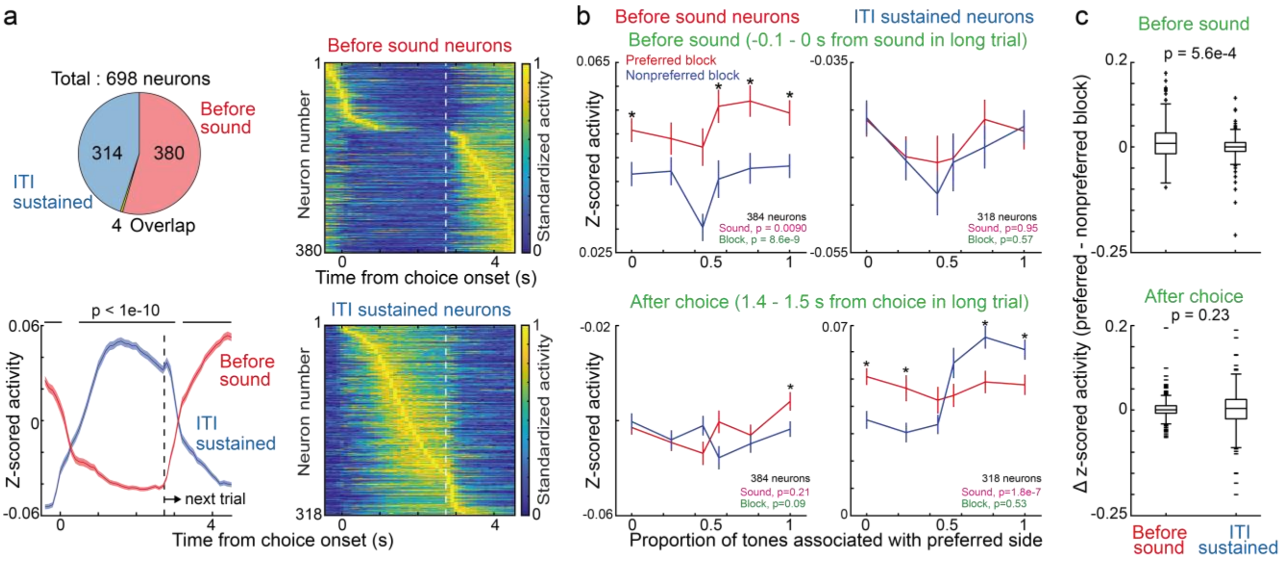


**Supplementary Fig. 6. Small overlapping of block modulated neurons and inter-trial-interval sustained neurons in the mPFC.**

**a.** Few overlapping between the neurons with increasing the activity before sounds (-0.1 – 0 s) (**Fig. 4b**) and those with sustained activity during inter trial interval (ITI). The sustained neurons had increased activity during 1 - 2 s after choice compared to the activity before the spout approaching (-0.2 - 0 s) ($p < 1e\text{-}10$, one-sided Wilcoxon signed rank test). Only 4 neurons were overlapped between the before-sound neurons and the ITI-sustained neurons (left top). The left bottom panel shows the means and standard errors of activity from choice onsets, suggesting the distinct activity patterns between the before-sound and ITI-sustained neurons (two-sided Mann-Whiteney U-test). Vertical dotted line shows the average timing of spout movement in the next trial. Right panels show the averaged and standardized activity of single neurons.

**b.** Activity of before-sound and ITI-sustained neurons in correct trials. Data presentations comply with **Fig. 4b**. Medians and robust standard errors (* $p < 0.01$ in the two-sided Wilcoxon signed rank test; top left: p = 0.0060, 0.026, 0.010, 3.0e-4, 9.0e-4, 7.6e-5; top right: p = 0.086, 0.88, 0.34, 0.82, 0.63, 0.85; bottom left: p = 0.16, 0.23, 0.21, 0.012, 0.078, 8.7e-6; bottom right: p = 1.6e-6, 1.7e-4, 0.018, 0.23, 0.0084, 0.0093).

**c.** Comparison of block modulations between the before-sound and ITI-sustained neurons. Before the sound onset, the block modulations were larger in the before-sound neurons than in the ITI-sustained neurons (two-sided Mann-Whitney U test) (central and edges of the box: median, 25th, and 75th percentiles; whiskers: most extreme data points not considered outliers (beyond 1.5 times the interquartile range)). Source data are provided as a Source Data file.

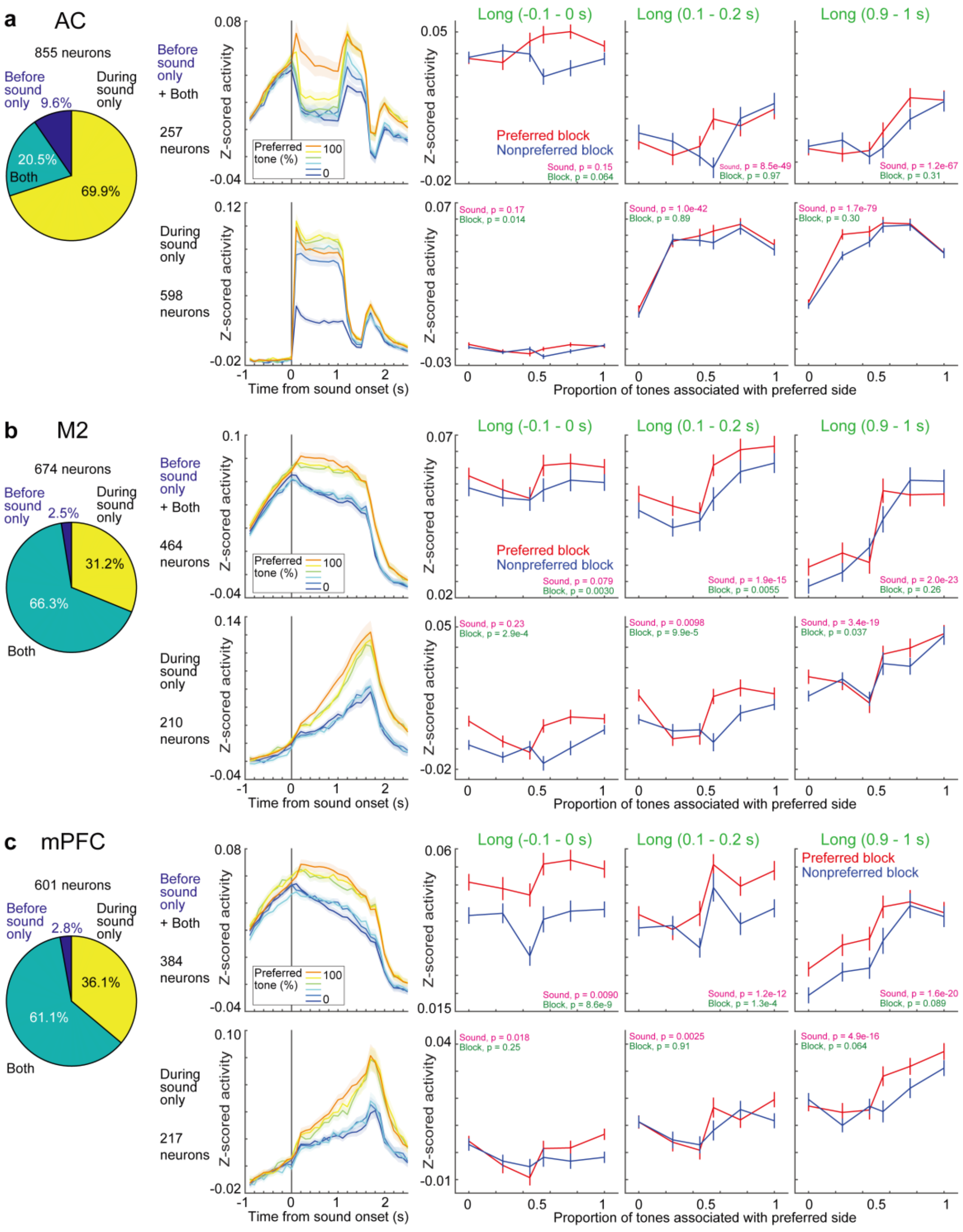


**Supplementary Fig. 7. Neurons started to activate before sounds modulate the neural representations from block to choice in the M2 and mPFC.**

We extracted 2 neuron groups from the task-relevant neurons. One group started to increase the activity from before sound onset in long-sound trials (-0.1–0 s) (defined as Before-sound-only or Both). The other group increased the activity only after sound onsets (During sound only). Pie charts show the distribution of neurons activated before, both, or during sounds. **a** – **c**. Top and bottom rows show the activity of neurons activated before- and during-sounds, respectively. In the M2 and mPFC, the before-sound neurons gradually changed the representations from block to choice. In contrast, the during-sound neurons mainly represented the choice. The population activity of AC had stable sound representations. P values in the two-way ANOVA are shown as the figure insets. Left: Means and standard errors. Right three figures: medians and robust standard errors. Data presentations comply with **Fig. 4 and 5**. Source data are provided as a Source Data file.

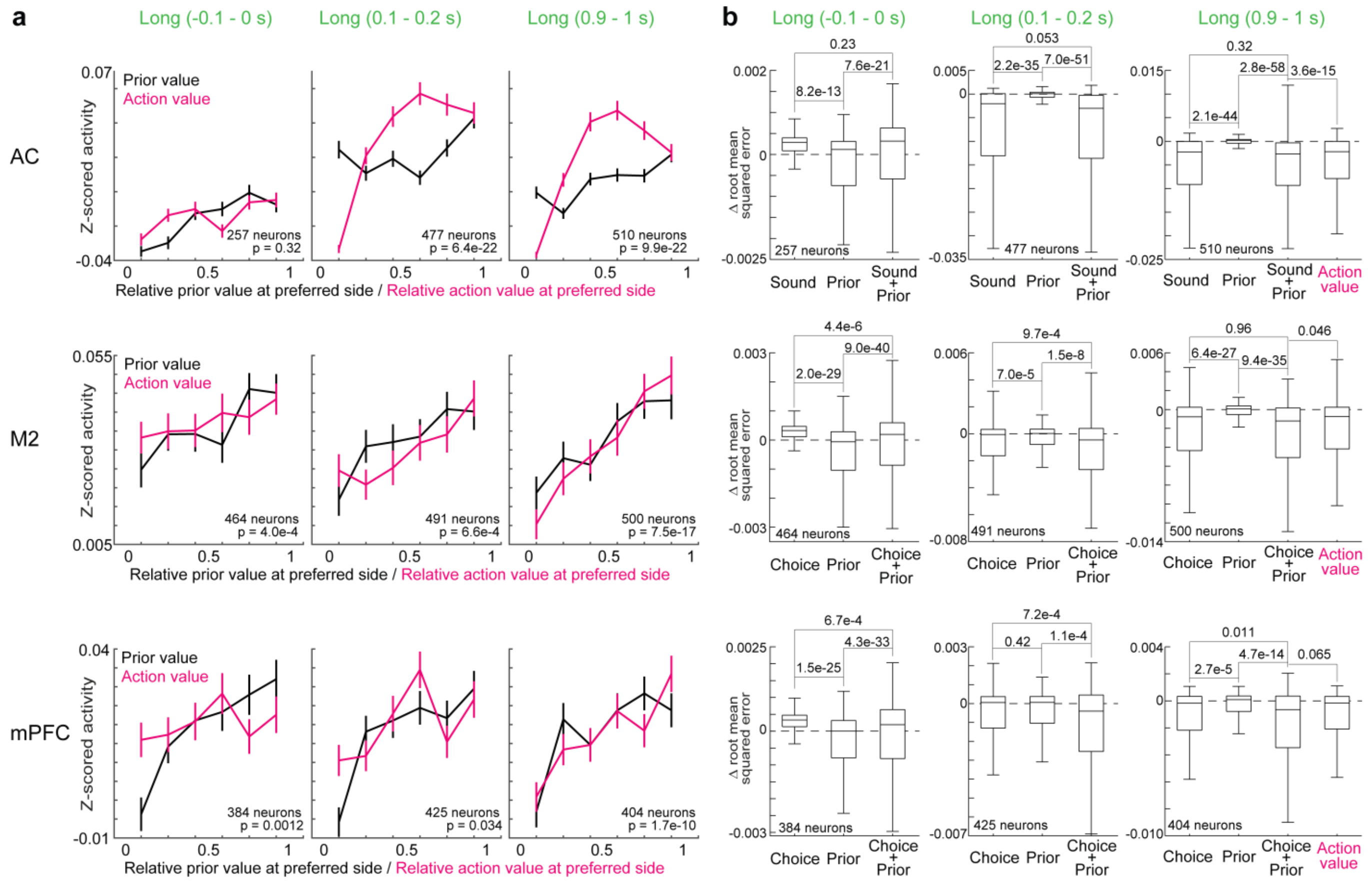


**Supplementary Fig. 8. Neural representations of task variables in different time windows before and during sound.**

**a.** Activity of task-relevant neurons in different prior values or action values. Data presentations comply with **Fig. 4d, 5i, and Supplementary Fig. 9i** but with action values. Medians and robust standard errors. We analyzed the Spearman partial correlations between the neural activity and prior values, or between the activity and action values without the effect of running speeds of mouse. Two-sided Wilcoxon signed rank test compared the correlation coefficients for the prior values and action values (p value in each inset). The prior and action values were correlated to the neural activity before and end of sounds, respectively, in the M2 and mPFC.

**b.** Neural encoding of task variables before and during sound. Linear regression with log-scaled neural activity analyzed which task parameters were represented in the neural activity of different time windows in long sound correct trials (**Methods**). We used 10-fold cross validation to test how the root mean squared error of the regression with only running speeds reduced by adding task parameters (Δ root mean squared error). In all the three regions, the prior values were represented before sound. In the AC and M2, the sound and choice were represented at the end of sounds, respectively. In the mPFC, the choice and prior values were additively represented. In the mPFC, there was no significant difference in the model fitting between the additive value (choice + prior) and action value (two-sided Wilcoxon signed rank test before Bonferroni correction, $p = 0.065$) (central and edges of the box: median, 25th, and 75th percentiles; whiskers: most extreme data points not considered outliers (beyond 1.5 times the interquartile range)). Box plots without the outliers. Source data are provided as a Source Data file.

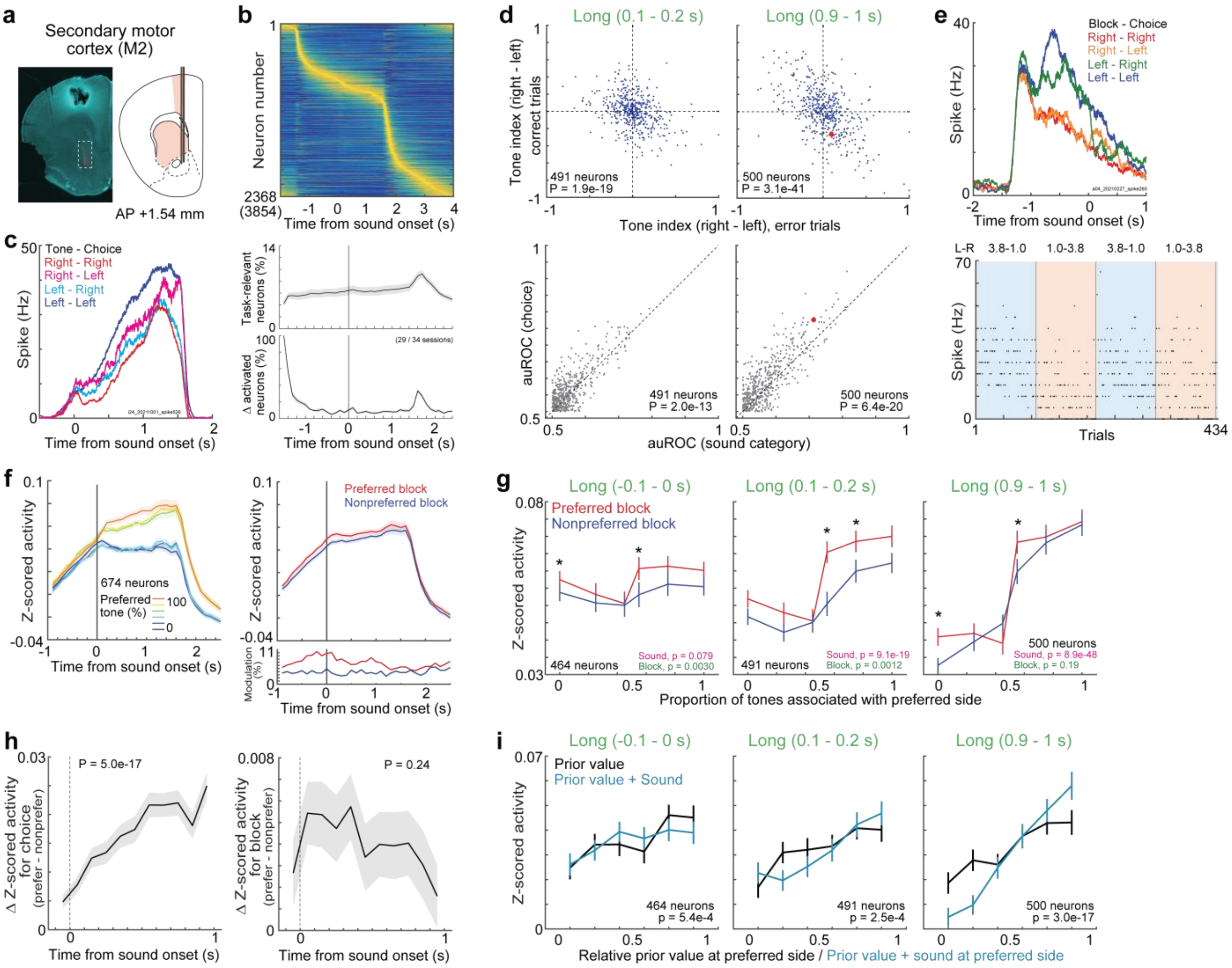


**Supplementary Fig. 9. Block and choice representations in the M2.**
Data are presented as in **Fig. 3, 4 and 5** for the M2 neurons.

**a.** Probe location for the M2 in an example mouse.
**b.** Average activity of task-relevant neurons in the M2. Bottom: Means and standard errors.
**c.** Average activity of an example neuron during long sounds.
**d.** Tone indices and area under the receiver operating curve (auROC) in task-relevant neurons (two-sided Wilcoxon signed rank test).
**e.** Average activity of an example neuron before sound onset.
**f.** Means and standard errors of activity in task-relevant neurons.

**g.** The activity was compared between sound categories and blocks with the two-way ANOVA (p values in the bottom right). Medians and robust standard errors (* $p < 0.01$ in the two-sided Wilcoxon signed rank test; -0.1–0s: p = 0.0018, 0.33, 0.17, 0.0033, 0.011, 0.043; 0.1–0.2s: p = 0.061, 0.15, 0.32, 8.6e-5, 5.0e-4, 0.13; 0.9–1s: p = 0.0088, 0.56, 0.064, 0.0057, 0.59, 0.30).

**h.** Sound- and block-dependent activity before and during sounds. Medians and robust standard errors (p values in the two-sided t-test in robust linear regression).

**i.** The neural activity was correlated to the prior values before sounds, while it was correlated to the prior value + sound (additive values) during sounds. Medians and robust standard errors. Two-sided Wilcoxon signed rank test compared the correlation coefficients for the prior values and additive values (p value in each inset). Source data are provided as a Source Data file.

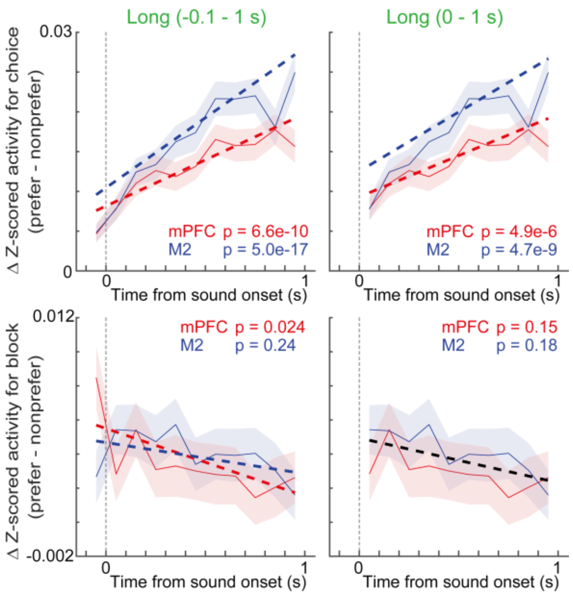


**Supplementary Fig. 10. Comparison of choice and block modulations of activity between the mPFC and M2.**
Data are same as in **Fig. 4c** and **Supplementary Fig. 9h** for the mPFC and M2. Robust linear regression showed that the choice modulations were significantly increased during sounds both in the mPFC and M2 (p values in insets, two-sided t-test). The block modulations were decreased in the mPFC. Medians and robust standard errors.

We then investigated whether the mPFC and M2 had different choice or block modulations with 10-fold cross validation (CV). We investigated whether the activity of mPFC and M2 fit to two independent robust linear regressions for mPFC and M2, or one regression by comparing the residual sum of squared error (RSS) between the two models. The CV separated the data into 10 groups. The 9 groups of data were used to train the robust regression. The regression was used to test the squared errors in 1 remaining group. We repeated this procedure 10 times to investigate the squared errors in all the data. To eliminate the noise from random grouping of CV, we repeated the CV for 1000 times and analyzed the average RSS.

We found that the choice modulations of mPFC and M2 fit to 2 independent regressions (average RSS: 29.71 and 29.00 for -0.1 – 1s and 0 – 1s) rather than 1 regression (29.78 and 29.06). The block modulation during -0.1 – 1 s were fit to 2 regressions (16.4416) compared to 1 regression (16.4422). In contrast, 1 regression was enough to model the block modulations of both the mPFC and M2 after sound onsets (0 – 1s: 14.6806 and 14.6795 for 2 and 1 regression). These results suggest that the choice modulation was larger in the M2 than mPFC. The block modulation was slightly larger in the mPFC than M2, but the difference was mainly observed before sound onset. Source data are provided as a Source Data file.

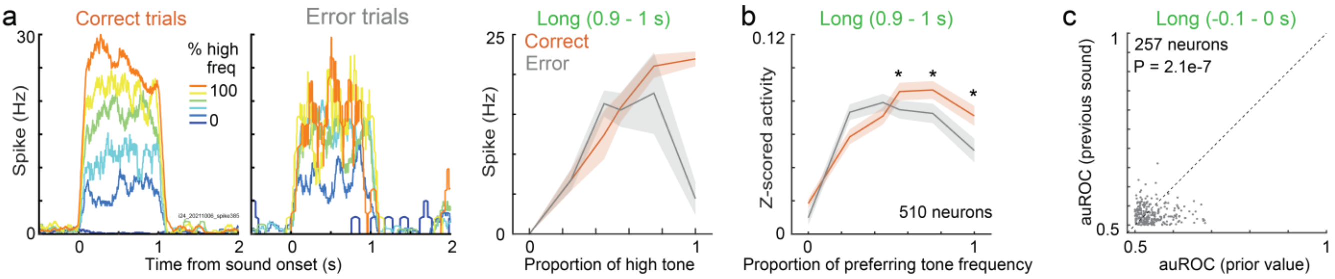


**Supplementary Fig. 11. Choice and prior-value modulation of neural activity in the auditory cortex (AC).**

**a.** Example sound-responsive neuron with choice modulation in the auditory cortex. (Left) Average neural activity during correct- and error-trials in long sound trials. (Right) Tuning curve. Means and standard errors.

**b.** Choice modulation in population auditory cortical activity. Medians and robust standard errors (*, $p < 0.001$ in the two-sided Wilcoxon signed rank test; p = 0.43, 0.69, 0.36, 4.0e-4, 3.4e-5, 2.0e-8).

**c.** Prior-value modulation of neural activity in the auditory cortex before sound. The area under the receiver operating curve (auROC) showed that the decoding accuracy of prior value was better than that of tone category of previous trial (two-sided Wilcoxon signed rank test). The neurons are same as **Supplementary Fig. 5b** before sound. Source data are provided as a Source Data file.

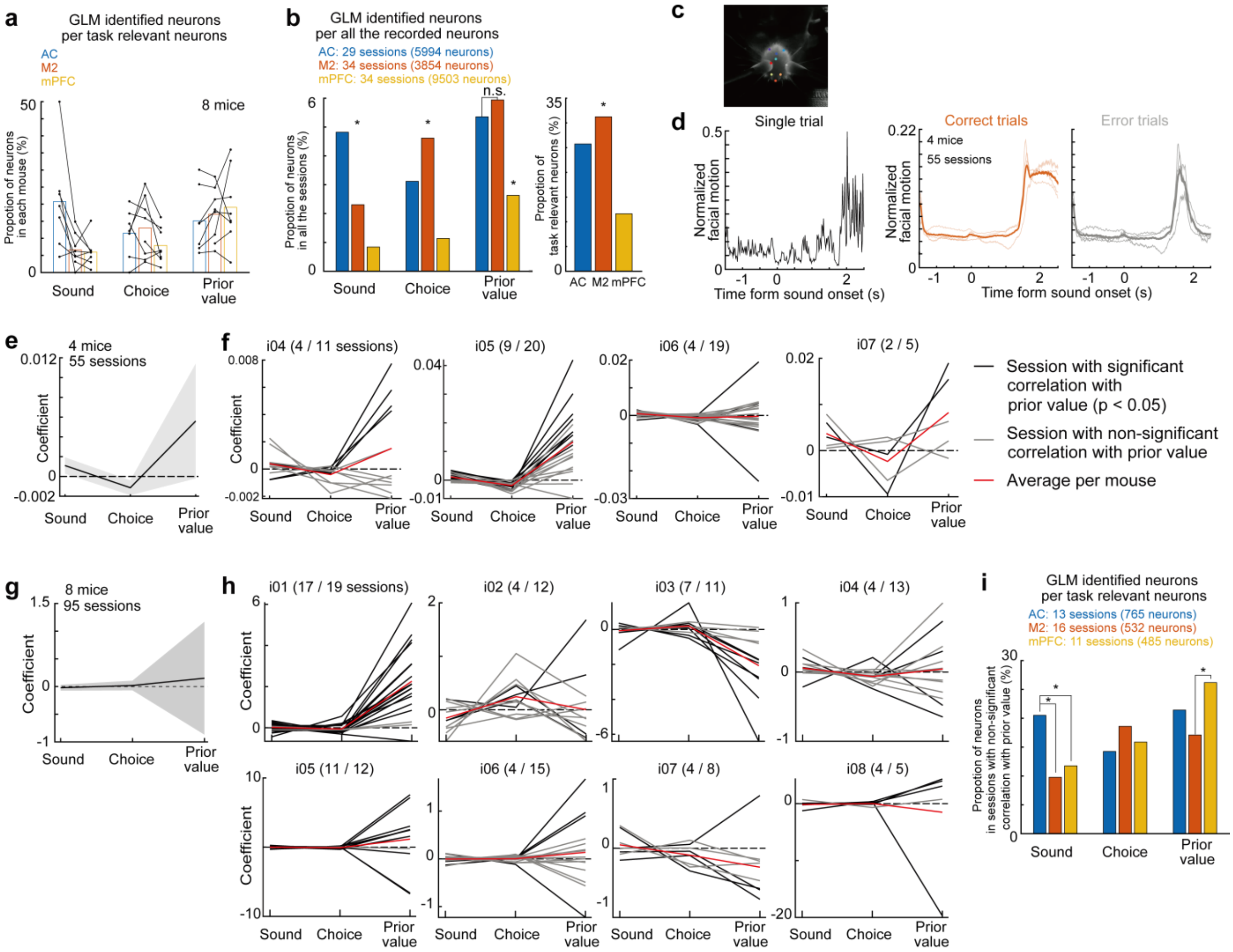


**Supplementary Fig. 12. Encoding of sounds, choices, and prior values in the AC, M2, and mPFC.**

**a.** Proportion of GLM-identified neurons per task-relevant neurons in each mouse. The GLM-identified neurons were detected as sound-, choice-, or prior-value-represented neurons. Bars show the average proportion of neurons across mice.

**b.** Proportion of GLM-identified neurons per all the recorded neurons. Data presentations comply with **Fig. 6c** but from all the recorded neurons. The proportion of neurons in mPFC decreased compared to **Fig. 6c**, because the proportion of task-relevant neurons in the mPFC was smaller than the other two regions (right). All the combinations except n.s. in the figure had significant differences in the proportions (* p < 0.01 in the two-sided chi-square test; left, sound, AC-M2, AC-mPFC, M2-mPFC, p = 2.4e-10, 2.1e-56, 6.2e-12; choice, p = 1.2e-4, 1.4e-18, 2.2e-36; prior value, p = 0.22, 1.8e-18, 1.1e-20; right, p = 2.9e-9, 3.3e-114, 3.3e-162).

**c.** Tracking of facial movement with DeepLabCut (DLC)[2]. We captured the facial movement of mouse with one camera with a frequency of 60 Hz (3 mice, 50 sessions) or 140 Hz (1 mouse, 5 sessions), as previous studies show that the neural activity and facial movements are correlated [3,4]. The camera was positioned below the spout to captured 9 face points (right/center/left/tip mouth, root/tip tongue, right/left/tip nose) and the spout movement. The spout movement was used to align the data from the video and the task events. We performed a linear completion when the data point had a likelihood below 0.6 as the post processing of

DLC. Missing data points were filled with the nearest available point except for the tongue movement.

**d.** Facial motion strength. In each of 9 facial features extracted by DLC, the velocities of features were defined as the absolute frame-by-frame difference of the xy position vector. Facial motion strength was defined as the sum of velocities in all the features. The data were scaled between 0 and 1 in each session. The trace of facial motion strength in single trial (left), and the average of all the mice and each mouse in correct (middle) or error trials (right) are shown.

**e.** Linear regression analyzed whether the tones, choices and prior values were correlated to the facial motion strength of mice. The facial motion strength was analyzed between the spout removal and the sound onset in each trial. Means and 95 % confidence intervals of regression coefficients (linear mixed-effects model; 55 sessions in 4 mice). The prior values did not have significant regression coefficients ($p = 0.059$ in the two-sided t-test).

**f.** Linear regression in each session. Red line shows the average regression coefficient per mouse. Parentheses show the number of sessions in which the prior values and facial motion strengths were correlated ($p < 0.05$ in the two-sided t-test).

**g.** Linear regression analyzed whether the tones, choices, and prior values were correlated to the running speeds of mice detected with rotary encoder. The running speed was analyzed between the spout removal and the sound onset. Means and 95 % confidence intervals of regression coefficients (two-sided t-test in linear mixed-effects model; 95 out of 97 sessions with neural recoding from the AC, M2, or mPFC, 8 mice; $p = 0.80$ for prior values). The remaining 2 sessions were excluded from the analysis, as the running speeds were not measured.

**h.** Data presentations comply with **f**, but for the running speeds of mice.

**i.** Data presentations comply with **Fig. 6c** but for the sessions in which the running speeds of mice were not significantly correlated with the prior values (gray sessions in **h**). The prior values were represented in all the three recorded regions (* $p < 0.01$ in the two-sided chi-square test; sound, AC-M2, AC-mPFC, M2-mPFC, p = 2.2e-7, 6.0e-5, 0.31; choice, p = 0.035, 0.43, 0.25; prior value, p = 0.054, 0.053, 4.2e-4). Source data are provided as a Source Data file.

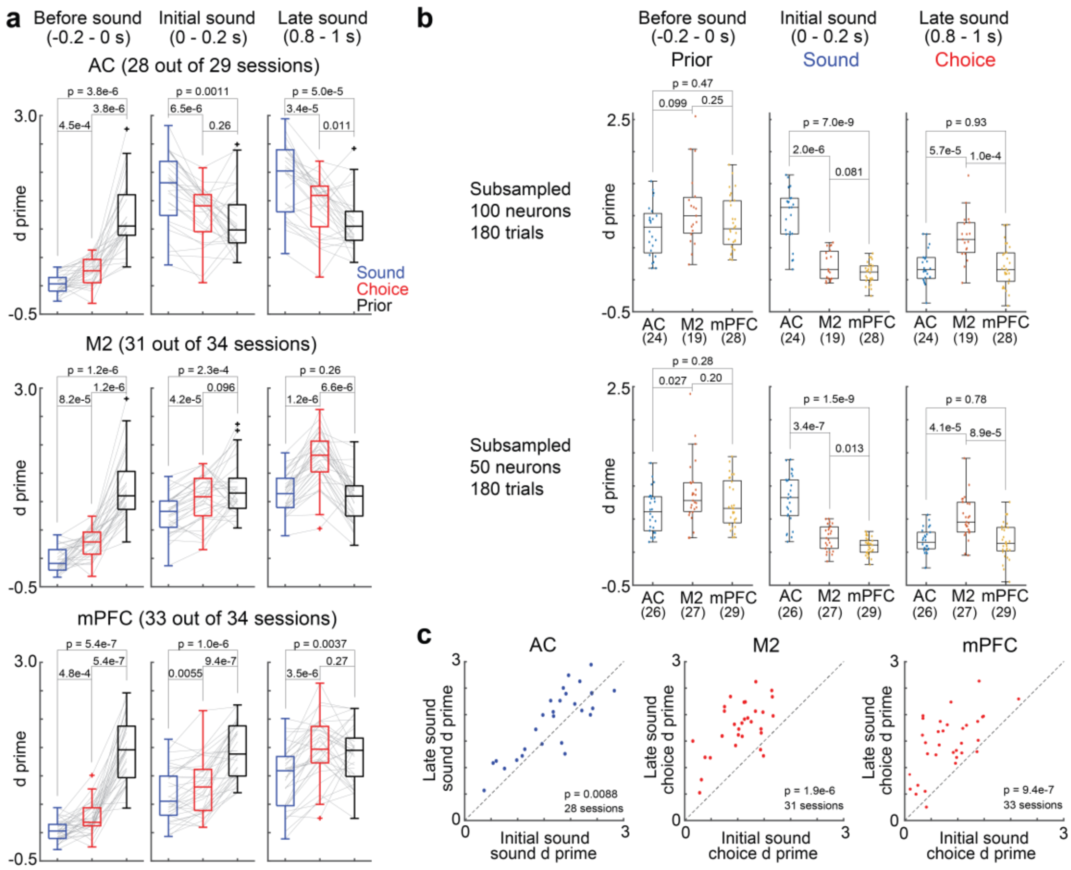


**Supplementary Fig. 13. Comparison of the decoding performance in the AC, M2, and mPFC with d prime.**

Data are presented as in **Fig. 7** but the decoding performance was plotted with d prime. **a.** Comparison of decoding performance within each cortical region (two-sided Wilcoxon signed rank test) (central and edges of the box: median, 25th, and 75th percentiles; whiskers: most extreme data points not considered outliers (beyond 1.5 times the interquartile range)).

**b.** Comparison across regions (two-sided Mann–Whitney U test) (central and edges of the box: median, 25th, and 75th percentiles; whiskers: most extreme data points not considered outliers (beyond 1.5 times the interquartile range)).

**c.** Comparison within each cortical region between initial and late sound phases (p value in the two-sided Wilcoxon signed rank test). Source data are provided as a Source Data file.

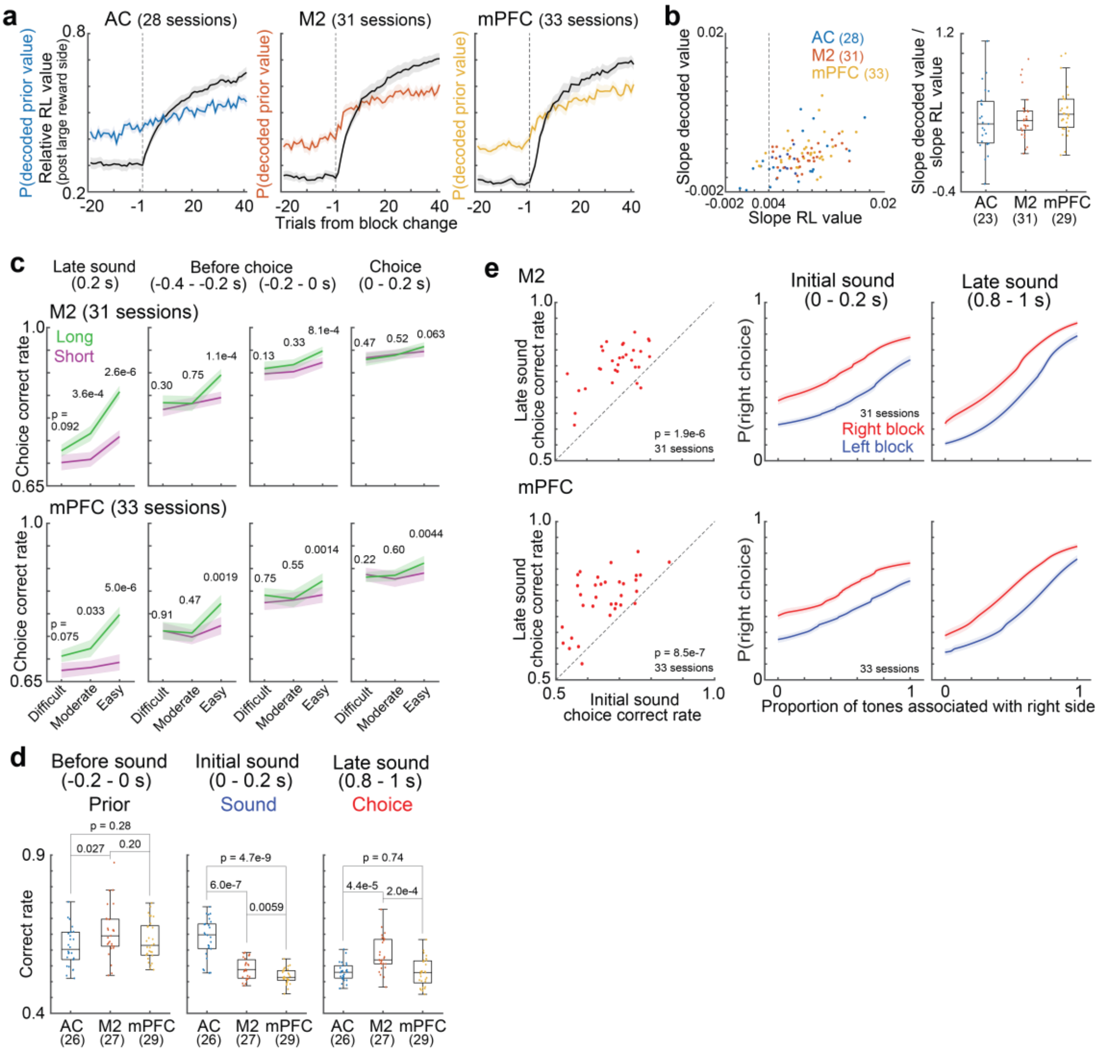


**Supplementary Fig. 14. Decoding performance in the AC, M2, and mPFC.**

**a.** Prior value decoding after the block changes. Relative RL value and P(decoded prior value) show the prior value in our RL model and the decoded prior value of high rewarded side after the block change, respectively. Both values were first averaged across the trials within a session and then averaged across the sessions. Error bars show the standard errors.

**b.** Updating of prior value decoding. We used the first 40 trials after the block changes and investigated the slope of RL prior value and that of decoded prior value in each session (regress in Matlab) (left). We used the sessions when the slope of RL value beyond 0.004 and investigated the ratio between the slopes of decoding and RL values (right). Parentheses show the number of sessions. Comparison of the slope ratio showed that the AC, M2, and mPFC did not have significant differences in the updating speed of prior value decoding (two-sided Mann–Whitney U test, p = 0.17, 0.20, and 0.65) (central and edges of the box: median, 25th, and 75th percentiles; whiskers: most extreme data points not considered outliers (beyond 1.5 times the interquartile range)).

**c.** Comparison of choice decoding performance between the long- and short-sound trials in the M2 and mPFC. Data are presented as in **Fig. 7c**, but categorized into different tone difficulties. Medians and robust standard errors (two-sided Wilcoxon signed rank test).

**d.** Comparison of the decoding performance in the AC, M2, and mPFC. Data are presented as in **Fig. 7d**, but 50 neurons and 180 trials were subsampled in each session for 10-fold cross validation (two-sided Mann–Whitney U test) (central and edges of the box: median, 25th, and 75th percentiles; whiskers: most extreme data points not considered outliers (beyond 1.5 times the interquartile range)).

**e.** Choice decoding in the mPFC and M2. Data are presented as in **Fig. 7e** but for choice decoding (two-sided Wilcoxon signed rank test). Means and standard errors. Source data are provided as a Source Data file.

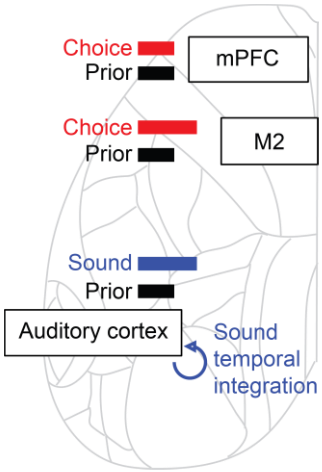


**Supplementary Fig. 15. Localized and global representation of prior value, sensory evidence, and choice in male mouse cerebral cortex.**
The mPFC neurons represented the additive combination of prior values and choices. The sounds and choices were selectively decoded from the auditory cortex and M2, respectively. These results suggest a localized computation of sound and choice which are required within single trials. All the recorded regions represented the prior values which needed to be maintained across trials. Our results suggest a localized and global computation of task variables essential for short- and long-time scales, respectively, in the cerebral cortex. The background image is the surface of mouse dorsal cortex based on the Allen Brain Atlas.
